# Temperature-induced changes in wheat phosphoproteome reveal temperature-regulated interconversion of phosphoforms

**DOI:** 10.1101/261065

**Authors:** Lam Dai Vu, Tingting Zhu, Inge Verstraeten, Brigitte van de Cotte, IWGSC, Kris Gevaert, Ive De Smet

**Affiliations:** Ghent University, Department of Plant Biotechnology and Bioinformatics, B-9052 Ghent, Belgium; VIB Center for Plant Systems Biology, B-9052 Ghent, Belgium; Department of Biochemistry, Ghent University, B-9000 Ghent, Belgium; VIB-UGent Center for Medical Biotechnology, B-9000 Ghent, Belgium; Lee’s Summit, Missouri, USA

**Keywords:** Wheat, Temperature, Phosphorylation, Signalling, Leaf, Spikelet, Phosphoproteomics

## Abstract

Wheat (*Triticum* ssp.) is one of the most important human food sources. However, this crop is very sensitive to temperature changes. Specifically, processes during wheat leaf, flower and seed development and photosynthesis, which all contribute to the yield of this crop, are affected by high temperature. While this has to some extent been investigated on physiological, developmental and molecular levels, very little is known about early signalling events associated with an increase in temperature. Phosphorylation-mediated signalling mechanisms, which are quick and dynamic, are associated with plant growth and development, also under abiotic stress conditions. Therefore, we probed the impact of a short-term increase in temperature on the wheat leaf and spikelet phosphoproteome. The resulting data set provides the scientific community with a first large-scale plant phosphoproteome under the control of higher ambient temperature, which will be valuable for future studies. Our analyses also revealed a core set of common proteins between leaf and spikelet, suggesting some level of conserved regulatory mechanisms. Furthermore, we observed temperature-regulated interconversion of phosphoforms, which likely impacts protein activity.

## INTRODUCTION

Wheat (*Triticum* ssp.) is one of the most important staple food crops around the world (Hawkesford *et al.*, 2013). However, the current production of wheat is predicted to be not sufficient to satisfy the future demands of the increasing world’s population (Hawkesford *et al.*, 2013; Mochida and Shinozaki, 2013; International Wheat Genome Sequencing Consortium (IWGSC), 2014). In addition, the global temperature is predicted to rise throughout the 21^st^ century (IPCC, 2014); and it has been estimated that for each degree (°C) of temperature increase, global wheat production will reduce by 6% impacting food security (Asseng *et al.*, 2015).

Wheat is sensitive to heat stress during all stages of its growth and development (Barber *et al.*, 2015; Akter and Rafiqul Islam, 2017). During wheat vegetative development, traits affected by high temperature include plant height, specific leaf weight, leaf width, relative water content, chlorophyll content and secondary metabolites (Akter and Rafiqul Islam, 2017). Furthermore, generative wheat growth and development are also very susceptible to increased temperatures (Bennett *et al.*, 1971; Sainiab *et al.*, 1983; Saini *et al.*, 1984; Draeger and Moore, 2017). Specifically, when wheat flowers are exposed to heat stress (10°C above the optimum condition) at the stage between ear initiation and anthesis (when anther development goes through meiosis) this causes abnormal development of the pollen grains in the anther and subsequently results in grain yield reduction (Sainiab *et al.*, 1983; Saini *et al.*, 1984; Fischer, 1985; Wardlaw *et al.*, 1989).

So far, transcriptome and proteome profiles were investigated in wheat under heat stress, revealing differences in gene expression and protein levels, respectively (Liu *et al.*, 2015; Wang *et al.*, 2016; Zhang *et al.*, 2017). Often, changes in gene expression for enzymes were in line with changes in the metabolite profiles upon stress (Rizhsky, 2004). Different metabolites, including organic acids, amino acids, polyols and lipidic compounds, which are beneficial for the plant during heat stress and known to protect the photosynthesis system, are enhanced in conditions of elevated temperature (Guy *et al.*, 2008; Scalabrin *et al.*, 2015; Qi *et al.*, 2017).

Several protein post-translational modifications (PTMs) are linked with plant stresses; but, these PTMs are hardly investigated in the context of temperature stress (Wu *et al.*, 2016; Hashiguchi and Komatsu, 2016). For example, protein phosphorylation is involved in the regulation of a large number of processes, including abiotic stress signalling (Kline *et al.*, 2010; Bonhomme *et al.*, 2012; Nguyen *et al.*, 2012; Zhang *et al.*, 2014*a*,*b*; Kanshin *et al.*, 2015). However, little is known about the phosphoproteome differences in the important crop wheat in vegetative and reproductive organs and during development under high temperature (Kumar *et al.*, 2017). Nevertheless, understanding PTM-mediated signalling cascades associated with an elevated temperature response is essential to gain insight in temperature tolerance and to facilitate future breeding (Rampitsch and Bykova, 2012).

Here, we monitored phosphorylation events in leaves of wheat seedlings and wheat spikelets exposed for 1h to higher temperature, and further analysed the data for biological processes potentially affected by phosphorylation. The information presented here not only improved our understanding about the role of protein phosphorylation in wheat under high temperature stress, but also provided a large number of phosphorylation sites for potentially critical proteins in this process. Furthermore, we observed temperature-regulated interconversion of phosphoforms, especially of neighbouring phosphosites, which likely impacts protein activity.

## MATERIAL AND METHODS

### Wheat Plant Materials and Growth Conditions

The seeds used in this study were from two bread wheat (*T. aestivum*, AABBDD, 2n = 6x = 42) cultivars, Fielder and Cadenza. The seeds were put on wet paper enclosed by plastic wrap and vernalized as such at 4°C for 3-4 days, and then transferred to room temperature for germination. Seeds that germinated uniformly were selected and grown in plastic pots containing soil at 21°C (Cadenza) or 24°C (Fielder) under 16 h light/8 h dark (100 μE m^−2^s^−1^ photosynthetically active radiation, supplied by cool-white fluorescent tungsten tubes, Osram), and 65–75% air humidity.

### Temperature Treatment

Temperature treatment was performed 8 h after the start of the light period. For the leaf material, Fielder plants at 7 days post germination growing in separate pots were transferred to two incubators and grown at 34°C (high temperature treatment) or 24°C (control temperature) under constant light (100 μE m^−2^s^−1^ photosynthetically active radiation) for 60 min. For the spikelet samples, Cadenza plants were cultivated in the greenhouse until the booting stage (stage 45 in Zadoks Decimal Code), then transferred to two incubators at respectively 34°C (high temperature treatment) and 21°C (control temperature) under constant light (100 μE m^−2^s^−1^ photosynthetically active radiation) for 60 min. The leaves of seedlings from Fielder and the spikelets in the middle section of the ears from Cadenza were collected and frozen in liquid nitrogen.

### qRT-PCR

Three biological replicates were used per time point. RNA was extracted and purified with the RNeasy Mini Kit (Qiagen) according to the manufacturer’s instruction for plant RNA extraction. DNA digestion was done on columns with RNase-free DNase I (Promega). The iScript cDNA Synthesis Kit (Biorad) was used for cDNA synthesis from 1 μg of RNA. qRT-PCR was performed on a LightCycler 480 (Roche Diagnostics) in 384-well plates with LightCycler 480 SYBR Green I Master (Roche) according to the manufacturer’s instructions. Two housekeeping genes, *ACTIN* (GenBank locus AB181991.1) and the *CELL DIVISION CONTROL PROTEIN* (*CDC*, GenBank locus Ta.46201) were used for normalization of the expression level of the *HEAT SHOCK PROTEIN*s. All the primers are listed in **Supplemental Table S1**.

### Protein Extraction and Phosphopeptide Enrichment

Total protein extraction was conducted on three biological replicate samples (leaf and spikelet material from independent plants) per wheat cultivar according to our previously described procedure with minor modifications (Vu *et al.*, 2017). Details can be found in the **Supplementary Information**. Phosphopeptides were enriched as previously described (Vu *et al.*, 2017).

### LC-MS/MS Analysis

Each sample was analysed via LC-MS/MS on an Ultimate 3000 RSLC nano LC (Thermo Fisher Scientific, Bremen, Germany) in-line connected to a Q Exactive mass spectrometer (Thermo Fisher Scientific). The peptides were first loaded on a trapping column (made in-house, 100 μm internal diameter (I.D.) × 20 mm, 5 μm beads C18 Reprosil-HD, Dr. Maisch, Ammerbuch-Entringen, Germany). After flushing the trapping column, peptides were loaded in solvent A (0.1% formic acid in water) on a reverse-phase column (made in-house, 75 µm I.D. x 250 mm, 1.9 µm Reprosil-Pur-basic-C18-HD beads, Dr. Maisch, packed in the needle) and eluted by an increase in solvent B (0.1% formic acid in acetonitrile) using a linear gradient from 2% solvent B to 55% solvent B in 120 min, followed by a washing step with 99% solvent B, all at a constant flow rate of 300 nl/min. The mass spectrometer was operated in data-dependent, positive ionization mode, automatically switching between MS and MS/MS acquisition for the 5 most abundant peaks in a given MS spectrum. The source voltage was set at 4.1 kV and the capillary temperature at 275°C. One MS1 scan (m/z 400−2,000, AGC target 3 × 10^6^ ions, maximum ion injection time 80 ms), acquired at a resolution of 70,000 (at 200 m/z), was followed by up to 5 tandem MS scans (resolution 17,500 at 200 m/z) of the most intense ions fulfilling predefined selection criteria (AGC target 5 × 10^4^ ions, maximum ion injection time 80 ms, isolation window 2 Da, fixed first mass 140 m/z, spectrum data type: centroid, under-fill ratio 2%, intensity threshold 1.3xE4, exclusion of unassigned, 1, 5-8, >8 positively charged precursors, peptide match preferred, exclude isotopes on, dynamic exclusion time 12 s). The HCD collision energy was set to 25% Normalized Collision Energy and the polydimethylcyclosiloxane background ion at 445.120025 Da was used for internal calibration (lock mass).

### Database Searching

MS/MS spectra were searched against the unpublished IWGSC RefSeq v1.0 database for *Triticum aestivum* (137052 entries) (wheat-urgi.versailles.inra.fr/Seq-Repository/Assemblies) with the MaxQuant software (version 1.5.4.1). For comparison, a second search against the earlier version of IWGSC PopSeq PGSB/MIPS v2.2 database (100344 entries), downloaded from wheatproteome.org, was performed. Detailed MaxQuant settings can be found in **Supplementary Information**. All MS proteomics data have been deposited to the ProteomeXchange Consortium via the PRIDE partner repository (Vizcaíno et al., 2014; Vizcaíno *et al.*, 2016) with the dataset identifier PXD008703. Next, the ‘Phospho(STY).txt’ output file generated by the MaxQuant search was loaded into the Perseus (version 1.5.5.3) data analysis software available in the MaxQuant package. Proteins that were quantified in at least two out of three replicates from each temperature were retained. Log2 protein ratios of the protein LFQ intensities were centered by subtracting the median of the entire set of protein ratios per sample. A two-sample test with a *p*-value cut-off *p*<0.01 was carried out to test for differences between the temperatures. Besides, phosphopeptides with 3 valid values in one condition and none in the other were also retained and designated “unique” for that condition.

### In Silico Analyses

For Gene Ontology (GO) analysis, the protein sequences of all identified phosphoproteins were loaded in the BLAST2GO software and blasted against the NCBI non-redundant protein sequence database of green plants (*Viridiplantae*) with a cut-off E-Value of 10^−5^. Afterwards, the results were examined for GO annotation and a Fisher’s exact test (p<0.05) was performed to extract enriched GO terms in the regulated phosphosite dataset. For Motif-X analyses, the Motif-X algorithm (Chou and Schwartz, 2011) was used to extract significantly enriched amino acid motifs surrounding the identified phosphosites. The sequence window was limited to 13 amino acids and foreground peptides were pre-aligned with the phosphosite in the centre of the sequence window. All identified proteins were used as the background dataset. The occurrence threshold was set at the minimum of 20 peptides and the P-value threshold was set at < 10^−6^. Structural modelling of the WD40 domain of *Ta*SPIRRIG was performed in SWISS-MODEL (Arnold *et al.*, 2006; Biasini *et al.*, 2014). The templates for the modelling studies were identified in the automated mode against the SWISS-MODEL template library (PDB: 5HYN). Structure representations were generated using the PyMOL Molecular Graphics System, Version 1.7.4, Schrödinger, LLC (www.pymol.org).

## RESULTS AND DISCUSSION

### Experimental Set-up for Early Leaf and Spikelet Phosphoproteome Analyses

So far, our knowledge on changes in the wheat proteome upon elevated temperature is largely limited to long-term exposures (day- or week-long treatments) (Majoul *et al.*, 2003; Laino *et al.*, 2010; Farooq *et al.*, 2011). We were interested in early signalling associated with a milder increase in ambient temperature, and therefore we wanted to profile changes in the phosphoproteome. To determine a suitable time point for proteome sampling, we first probed the expression levels of two *HEAT SHOCK PROTEINs*, since early thermal sensing is largely reflected in the transcription of *HEAT SHOCK PROTEINs* (Xu *et al.*, 2011). Here, we exposed 7 days old wheat seedlings (Fielder) grown at 24°C for a short-term treatment of 34°C and harvested whole shoots at different incubation times (**Figure 1A**). Recent evidence in cereal crop plants has demonstrated a link between high temperature sensitivity at booting stage and seed yield (Hedhly *et al.*, 2009; Draeger and Moore, 2017). Hence, we used booting wheat plants (Cadenza) grown at 21°C and exposed to increased ambient temperature (34°C), after which we harvested spikelets at different incubation times (**Figure 1B**). Since developmental stages differ in optimal growth temperature (Porter and Gawith, 1999), we chose different optimal growth temperatures as the control condition for our experiment. We analysed the transcription of *TaHSP70d* and *TaHSP90.1*, which are markers for temperature response (Xue *et al.*, 2014), in both leaf and spikelet samples. We found that the transcriptional response of *TaHSP70d* and *TaHSP90.1* peaks in both samples at 60 min, indicating a maximum of early high temperature response (**Figure 1C-D**). Therefore, to identify early phosphorylation-controlled signalling components associated with a mild increased temperature in wheat, we subjected both leaf and spikelet samples from the 60 min time point to our phosphoproteomic workflow (Vu *et al.*, 2016).

**Figure 1.**
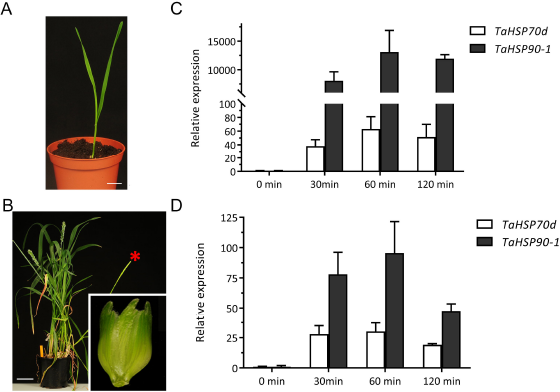
Different wheat cultivars and organs used in this study. **(A)** Fielder seedlings are depicted at 7 days after germination. Scale bar, 2.2 cm. **(B)** Cadenza spikelet (inset) is depicted from plants at the booting stage. Red asterisk indicates representative ear used for sampling. Scale bar, 7.5 cm. **(C-D)** Analysis of *HSP70* and *HSP90* expression in both leaf and ear as a proxy for the heat sensing shows a maximum increase at 60 min after transferring to high temperature.

### New wheat reference sequence improves protein identification

Advances in the wheat reference sequence assembly provide a solid basis for proteome studies in wheat (Brenchley *et al.*, 2012; International Wheat Genome Sequencing Consortium (IWGSC), 2014; Luo *et al.*, 2017). Through Ti-IMAC enrichment and subsequent LC-MS/MS analysis, we identified 3822 phosphopeptides containing 5178 phosphorylated amino acids, representing 2213 phosphoproteins in the leaf samples using the unpublished IWGSC RefSeq v1.0 assembly (**Figure 2 and Supplementary Table S2**). In spikelet samples, our workflow led to the identification of 5581 phosphopeptides containing 7023 phosphosites located on 2696 proteins (**Figure 2 and Supplementary Table S3**). As a comparison, we performed a second search using the earlier published protein sequence database based on the draft genome sequences of bread wheat (International Wheat Genome Sequencing Consortium (IWGSC), 2014). The new protein database, based on the unpublished IWGSC RefSeq v1.0 assembly, resulted in an increase of 30% and 34% of identifications compared to the search using the previous search database that identified 3975 and 5234 phosphosites for leaf and spikelet samples, respectively. This seems to correlate with the increase of 36.5% in the number of entries in the new database compared to the old database, supporting the quality of the new wheat reference sequence assembly. To our knowledge, this is currently the largest set of identified phosphosites in the *Triticum* family. The identified phosphosites in this study were added to our PTMViewer (bioinformatics.psb.ugent.be/webtools/ptm_viewer/) (Vu *et al.*, 2016). In addition, we found several phosphosites that were differentially regulated between normal (21 or 24°C) and increased ambient temperature (34°C) in wheat leaves and spikelets (**Figure 2).**

**Figure 2.**
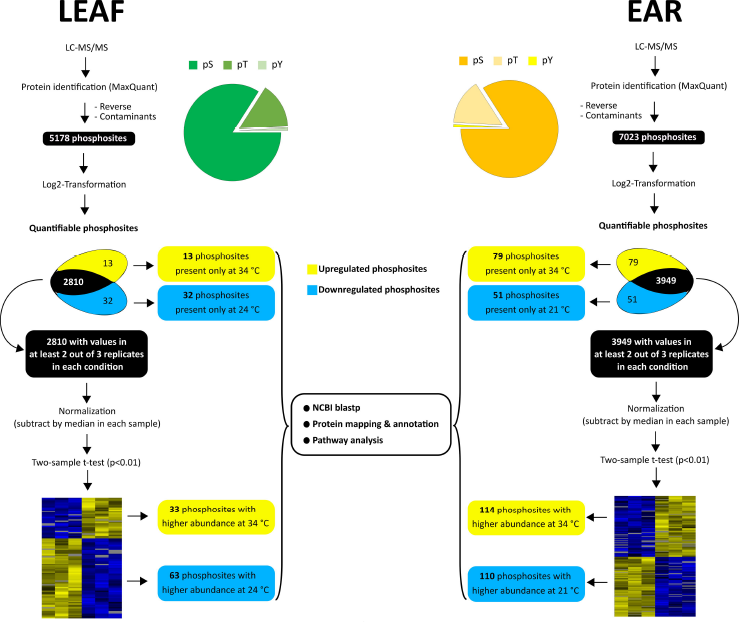
Summary of the phosphoproteome analysis in wheat leaf and ear. T-test significant hits and phosphosites with valid values reproducibly present in only one condition in each organ are collectively analyzed and called as upregulated or downregulated phosphosites.

### A Temperature-regulated Wheat Leaf Phosphoproteome

Phosphosites that exhibited valid values in one condition and none in the other indicate a massive change in phosphorylation levels. For the wheat leaves, we could identify 13 phosphosites that only occurred in the 34°C samples and 32 phosphosites unique for the 24°C condition (**Figure 2 and Supplementary Table S4**). On the rest of the wheat leaf dataset, we performed a Student’s t-test (p<0.01) on phosphosites with at least 2 valid values in any condition (2810 phosphosites), and this resulted in 33 significantly upregulated phosphosites and 63 significantly downregulated phosphosites at high temperature (**Supplementary Table S5**). Proteins with phosphosites uniquely identified in either conditions and significantly deregulated phosphoproteins from the statistical test were combined and analysed for overrepresented GO terms in biological processes (**Figure 3**) and molecular function (**Supplementary Figure S2**). As expected, upregulated phosphoproteins are highly enriched in the GO terms of stress-induced processes such as response to heat, protein folding (Zhu, 2016), response to hydrogen peroxide (Gupta *et al.*, 2016) and glucose transport (Ruan *et al.*, 2010). On the other hand, downregulated phosphoproteins were mainly enriched in positive regulation of translational elongation/termination, and ribosome biogenesis (Cherkasov *et al.*, 2015).

**Figure 3.**
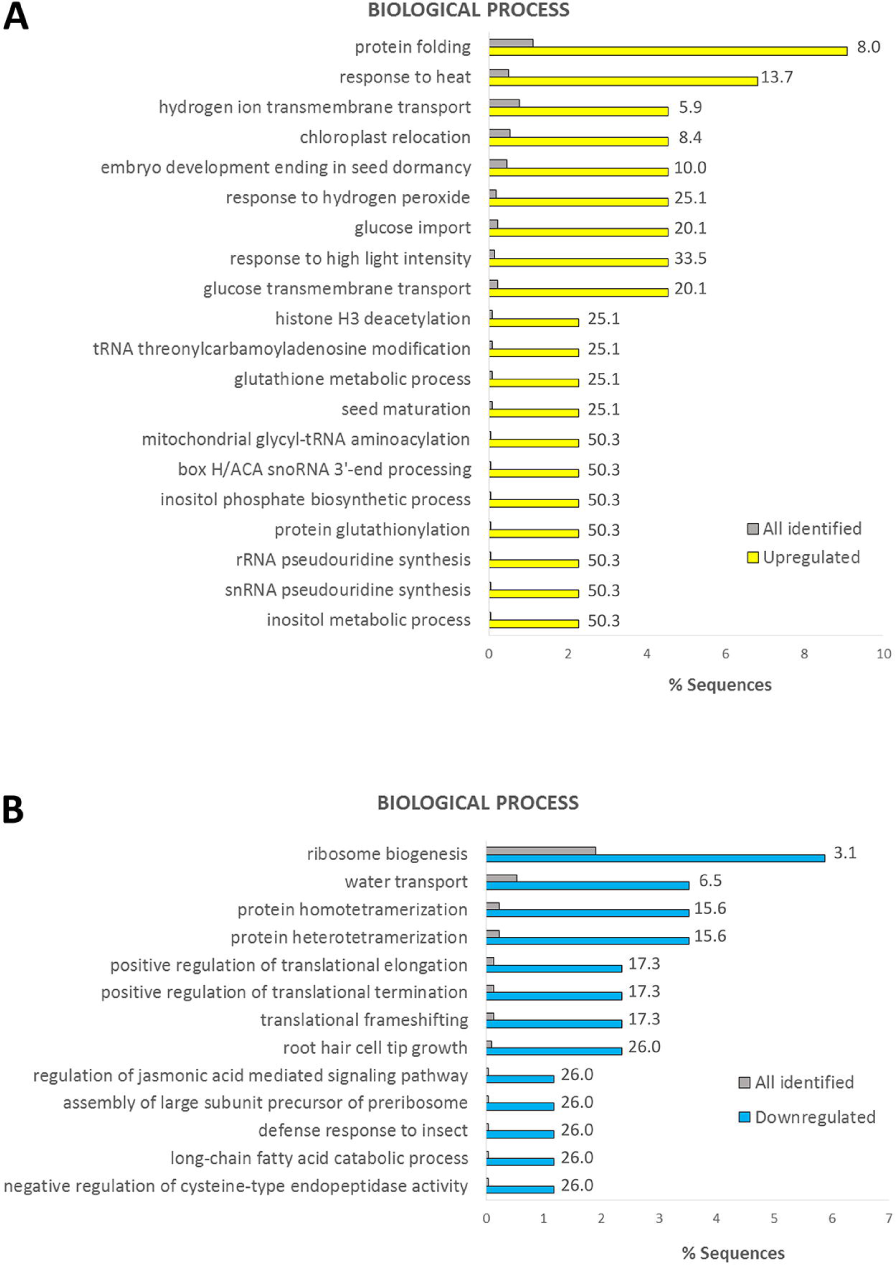
GO enrichment for biological process in upregulated (A) and downregulated (B) phosphoproteins in leaf samples. All identified leaf phosphosites were used as the background dataset. Fold change is indicated.

Expression of *HSPs* is rapidly induced in the leaf by increased temperature (**Figure 1C**), as the resulting proteins play crucial roles when plants are exposed to increased temperature (Sun *et al.*, 2002; Kotak *et al.*, 2007). In our leaf data set, we furthermore identified several differential phosphorylation sites of HSPs at 34°C (**Supplementary Table S4-5**), namely HSP90 (TraesCS2A01G033700.1, *Ta*HSP90) and HSP60-3A (TraesCSU01G009200.1, *Ta*HSP60-3A) were 10.4-fold and 4.6-fold upregulated at S224 and S577, respectively. However, for both proteins, another phosphosite, namely S93 of *Ta*HSP90 and T420 of *Ta*HSP60-3A, was not differentially phosphorylated after 1 h exposure to 34 °C. This suggested that HSP90 and HSP60-3A protein abundance is likely not the basis for the increase in S224 and S577 phosphopeptide increase, respectively.

Noticeably, our dataset indicated that the phosphoproteome of the photosynthesis machinery in wheat leaves is severely affected by high temperature (**Supplementary Tables S4 and S5**). For example, phosphorylation of T33, T37 and T39 of the subunit P of photosystem I (TraesCS2A01G235000.1) was 3.2-fold downregulated after 1 h exposure to 34°C (**Supplementary Table S5**). In addition, an actin-binding protein (TraesCS1D01G422700.2), whose homologue in *Arabidopsis* (CHUP1) is important for proper chloroplast positioning (Oikawa *et al.*, 2008), was found to be considerably less phosphorylated at S157 upon high temperature (**Supplementary Table S4**). Besides, a kinesin-like protein (TraesCS7D01G176200.1, homologous to *Arabidopsis* KAC1) is highly phosphorylated in its kinesin motor domain (S444) in response to high temperature (**Supplementary Table S4**). Both CHUP1 and KAC1 regulate the accumulation of chloroplast actin filaments in *Arabidopsis*, thus facilitating the anchorage of chloroplasts on the plasma membrane. Last, phosphorylation of kinases involved in chloroplast movement such as the phototropin homologues TraesCS5D01G389200.2 and TraesCS2B01G290500.3 (S525 and S294, respectively) was also elevated by heat (**Supplementary Table S4 and S5**).

The post-translational import of chloroplast proteins is a highly regulated process (Strittmatter *et al.*, 2010). Our dataset shows several components of this process to be affected by high temperature. Increased temperature also highly induced the phosphorylation of a wheat homologue (TraesCS5D01G132600.1) of *Arabidopsis* STY46 kinase at S31 (**Supplementary Table S4**). In *Arabidopsis*, STY46 and its homologues STY8 and STY17 facilitate import of chloroplast preproteins by phosphorylation of their N-terminal transit peptide (Lamberti *et al.*, 2011). On the other hand, many chloroplast proteins are integrated into the chloroplast outer membrane (COM) without any cleavable signal sequence (Hofmann and Theg, 2005). The ANKYRIN REPEAT-CONTAINING PROTEIN 2 (AKR2) interacts with chloroplast specific lipid markers and facilitates the insertion of COM proteins into the chloroplast outer membrane (Kim *et al.*, 2014). It is speculated that the regulatory mechanism of this process involves conformational changes of AKR2 via PTMs (Kim *et al.*, 2014). Here, we showed that phosphorylation of the AKR2 homologue in wheat (TraesCS4A01G328600.1) at S404 is two-fold upregulated in response to higher temperature (**Supplementary Table S5**). While protein import in chloroplasts has been shown to be altered under stress conditions (Dutta *et al.*, 2009; Ling and Jarvis, 2016), our dataset indicated that this response, especially to high temperature, is highly regulated by phosphorylation.

In conclusion, our temperature-mediated leaf phosphoproteome pinpointed photosynthesis as a central target of higher temperature and identified several phosphorylated residues on key components for further functional characterization.

### A Temperature-regulated Wheat Spikelet Phosphoproteome

For the wheat spikelet, we identified 79 phosphosites that are only present in the 34°C samples and 51 phosphosites that are unique for 21°C samples (**Figure 2 and Supplementary Table S6**). A Student’s t-test (p<0.01) was performed on the rest of the wheat spikelet dataset (phosphosites with at least 2 valid values in temperature condition; 3949 phosphosites), and this resulted in 114 significantly upregulated phosphosites and 110 significantly downregulated phosphosites at elevated temperature (**Supplementary Table S7**). Proteins with phosphosites uniquely identified in either condition and significantly deregulated phosphoproteins from the statistical test were combined and GO analysis was performed similarly as for the leaf samples (**Figure 4 and Supplementary Figure S3**). The biological processes enriched in leaf samples were also increased here, such as protein folding, response to heat, response to hydrogen peroxide. Similar to the leaf GO enrichment (**Figure 3**), terms associated with translation were predominantly enriched for downregulated phosphoproteins.

**Figure 4.**
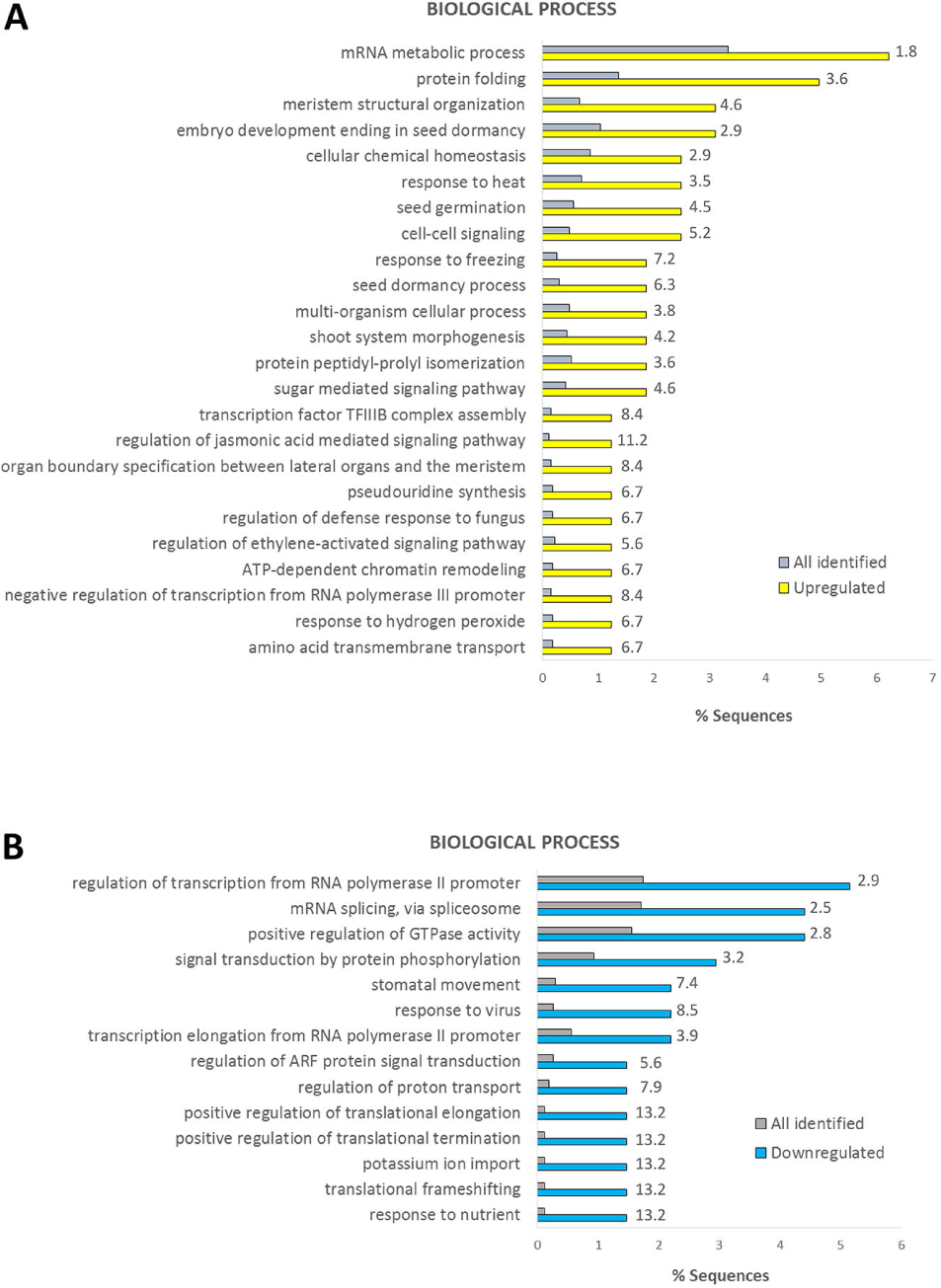
GO enrichment for biological process in upregulated (A) and downregulated (B) phosphoproteins in ear samples. All identified ear phosphosites were used as the background dataset. Fold change is indicated.

Reproductive development in plants is known to be greatly dependent on the epigenetic control of expression of flowering genes (Gan *et al.*, 2013). This often involves histone modifications such as (de)acetylation, methylation and ubiquitination (Lawrence *et al.*, 2016). Here, we found that the phosphorylation of several histone-modifying enzymes was deregulated in response to heat. For example, the phosphoserine 297 of the histone deacetylase TraesCS1A01G445700.3 was 7.1-fold downregulated and the phosphorylation of S762 S763 in the histone-lysine N-methyltransferase TraesCS2A01G262600.1 was 2.4-fold decreased (**Supplementary Table S7**). In contrast, an ubiquitin protease, TraesCS4D01G266600.3, was 2.2-fold more phosphorylated at S31 and T32. Its *Arabidopsis* homologue, *UBP26*, deubiquitinates the histone H2B to regulate floral transition by control the expression of *FLOWERING LOCUS C* (*FLC*) (Schmitz *et al.*, 2008). Furthermore, it has been demonstrated that phosphorylation is crucial for the activity of histone-modifying enzymes (Pflum *et al.*, 2001; Schmitz *et al.*, 2008; Xu *et al.*, 2015).

Another important step in epigenetic control of gene expression is the ATP-dependent restructuring of nucleosomes (Vignali *et al.*, 2000). Phosphorylation of two homologous SWI2/SNF2 class of chromatin remodelling ATPases, TraesCS7D01G206700.3 (at T2492) and TraesCS7B01G110600.1 (at S1668 and S1671), was massively induced by heat (**Supplementary Table S6**). The *Arabidopsis* homologue, SPLAYED (SYD), is known to be a co-repressor during floral transition (Wagner and Meyerowitz, 2002). In contrast, phosphorylation of S1728 in the SNF2 ATPase TraesCS6B01G048200.2 is 1.7-fold downregulated (**Supplementary Table S7**). Its homologue in *Arabidopsis*, BRAHMA (BRM) plays a pivotal role in controlling flowering time by regulating the expression of *FLC* and inflorescence architecture, mainly via interaction with the transcription factor KNAT1 (Zhao *et al.*, 2015). Interestingly, a wheat homologue of KNAT1, TraesCS5B01G410600.1, was also less phosphorylated at high temperature (**Supplementary Table S7**).

In conclusion, our data suggested that an increase in ambient temperature can alter phosphorylation status of chromatin remodelling proteins as an important mechanism to control gene expression during the reproductive stage. Further, other proteins involved in pollen, pistil or gametophyte development (**Supplementary Tables S6 and S7**) also exhibit altered phosphorylation in response to increased temperature.

### Comparison of Leaf and Spikelet Phosphoproteome

In total, we identified 2491 identical phosphosites in both organs, which account for 48% and 35% of all identified phosphosites in leaf and spikelet samples, respectively (**Figure 5**). Only 7 phosphosites were found commonly upregulated at high temperature in both organs and 8 were commonly downregulated in both organs (**Figure 5 and Supplementary Table S8**). Notwithstanding the considerable overlap between the phosphosites identified in both organs, the limited overlap between similarly regulated phosphosites indicated distinct responses in the leaf and spikelet phosphoproteomes at the early stages of thermal signalling.

**Figure 5.**
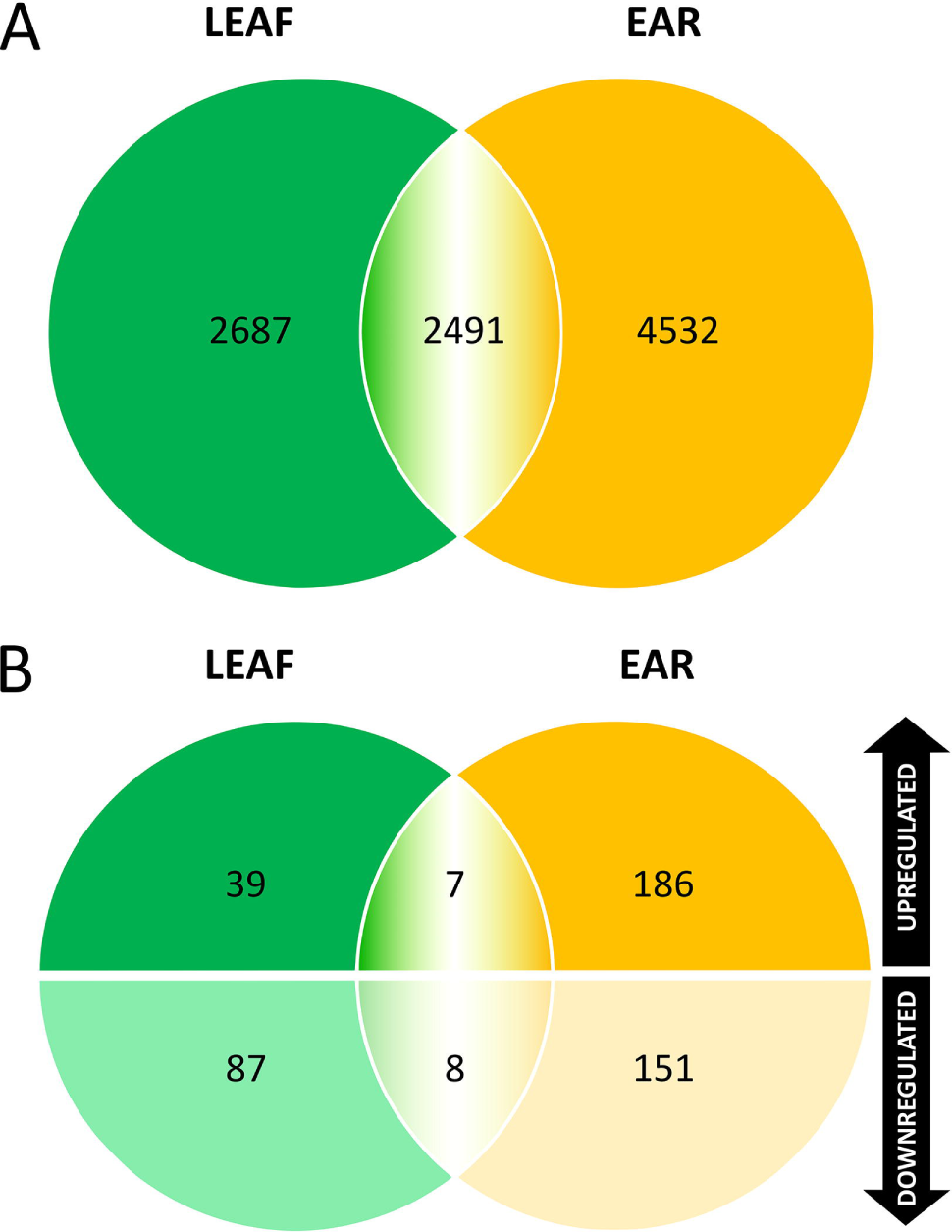
Venn diagrams showing the number of common identified phosphosites as well as deregulated phosphosites in leaf and ear samples.

Among the common higher temperature-induced phosphosites, phosphorylation of S464 of the pseudouridine synthase TraesCS2B01G177000.1 was increased 1.6-fold and 1.9-fold in leaf and spikelet samples, respectively. Pseudouridylation of mRNA as well as of non-coding RNAs can be induced in stress conditions and is important for the regulation of gene expression, and involved in splicing, translation and decay of mRNA (Karijolich *et al.*, 2015). On the other hand, three different translation initiation factors are present among the commonly regulated proteins with downregulated phosphosites (**Supplementary Table S8)**. This is in agreement with heat stress-triggered overall pausing of translation elongation, and with heat-induced HSP70 protecting cells from heat shock-induced pausing (Shalgi *et al.*, 2014; Merret *et al.*, 2015). Especially, dephosphorylation of translation initiation factors correlates with the reprogramming of translation following thermal stress in wheat (Gallie *et al.*, 1997).

### Leaf and Spikelet Phosphoproteome Motif-X Analyses Reveal Distinct Regulation of Phosphorylation Motifs

So far, little is known about the protein kinases and phosphatases involved in temperature signalling (Ding *et al.*, 2015; Yu *et al.*, 2017; Li *et al.*, 2017; Zhao *et al.*, 2017). Therefore, we used the identified phosphosites to reveal potential phosphorylation motifs and associated kinases that may act in a high-temperature responsive manner. The Motif-X algorithm was applied on the set of regulated phosphosites in leaf and spikelet samples separately, using the sequences of all identified phosphoproteins in either organ as reference (**Figure 6**). In the spikelet, the common SP motif was enriched in both upregulated as well as downregulated phosphosites. This suggested that kinases (and phosphatases) targeting those sites are tightly regulating the protein phosphorylation signatures (meaning the specific combination of phosphorylated and non-phosphorylated residues), which impacts on overall protein behaviour, such as protein activity and localization (Salazar and Höfer, 2009). The acidic SD motif was significantly overrepresented among the upregulated phosphosites (3.61-fold). In contrast, the downregulated phosphosites showed an enrichment in the basic RxxS motif (4.37-fold) (**Figure 6**). This latter trend was also found in the leaf samples (**Figure 6**). Despite that no motif enrichment was obtained for the upregulated phosphosites in leaf, due to the small size of the data set, we identified six SD motifs among these sites, which account for 13% of the upregulated phosphosites in leaves. This was comparable with 14% of the upregulated phosphosites in the spikelet samples which also shows the SD motif. This possibly indicated a common molecular mechanism of higher temperature response via phosphorylation across different organs and different growth stages. While local intracellular parameters such as the pH can slightly vary in a temperature-dependent manner and thus affect the property of amino acid residues around the phosphosites (Wilkinson, 1999; Schönichen *et al.*, 2013), we do not rule out the possibility that certain phosphosites are targeted by a specific set of higher temperature-activated kinases. The acidic motif SD is known to be targeted by MAP kinases (MPKs), receptor-like kinases (RLKs) and calcium-dependent protein kinases (CDPKs), while RxxS is a motif commonly targeted by MAP kinase kinases (M2Ks) (van Wijk *et al.*, 2014). In support of this, we found 6 RLKs among 10 kinases with a higher phosphorylation level at 34°C in the ear, whereas 3 out of 7 kinases with decreased phosphorylation level are predicted to have MAP3K or MAP4K activity **(Supplementary Table S9)**.

**Figure 6.**
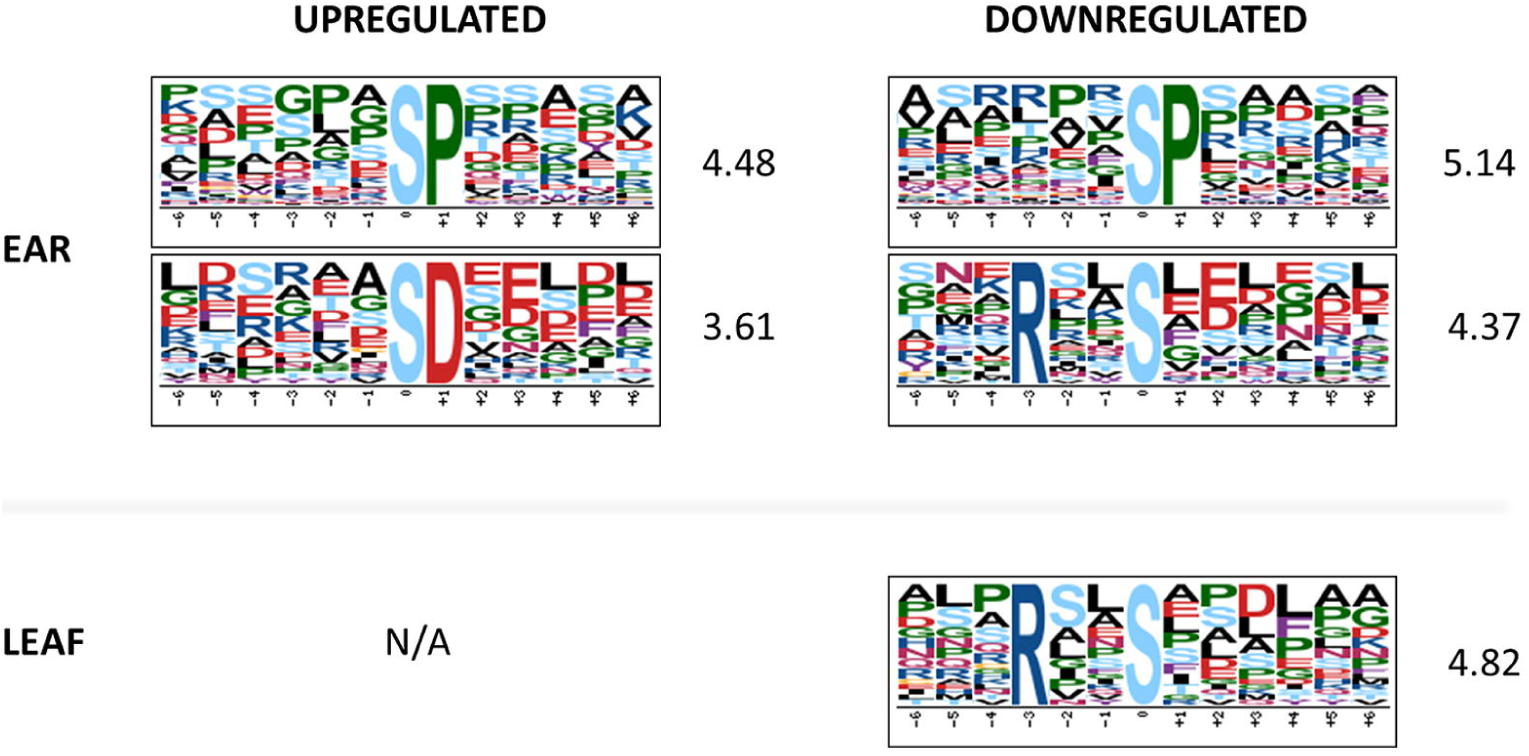
Motif-X analysis show an enrichment of an acidic phosphomotif among upregulated phosphosites and of a basic motif among downregulated phosphosites in leaf and ear. Fold-change of the enrichment compared to the background dataset are indicated. N/A, not available.

#### Phosphoproteins with multiple deregulated phosphosites

Since the protein phosphosignature will determine protein behaviour (Salazar and Höfer, 2009), we probed the leaf and spikelet phosphoproteome data for proteins that displayed a combination of up and down-regulated phosphosites. We found 13 phosphoproteins in the spikelet samples and one in the leaf samples that contained both significantly up and downregulated phosphosites (**Table 1**). It is thus very likely that the status of these phosphosites is not affected by changes in the protein level, but rather by higher temperature-dependent activity of associated kinases and phosphatases. These protein phosphatases and kinases might be activated by higher temperature and target the phosphosites independently to generate different phosphoforms of the target protein (**Figure 7A**). However, the phosphorylation and dephosphorylation events might also occur in an interdependent manner upon higher temperature (**Figure 7B**) (Salazar and Höfer, 2009; Nishi *et al.*, 2015). Crosstalk between different or the same type of PTMs is very common (Beltrao *et al.*, 2013; Nishi *et al.*, 2015), but is still not widely explored in plants.

**Figure 7.**
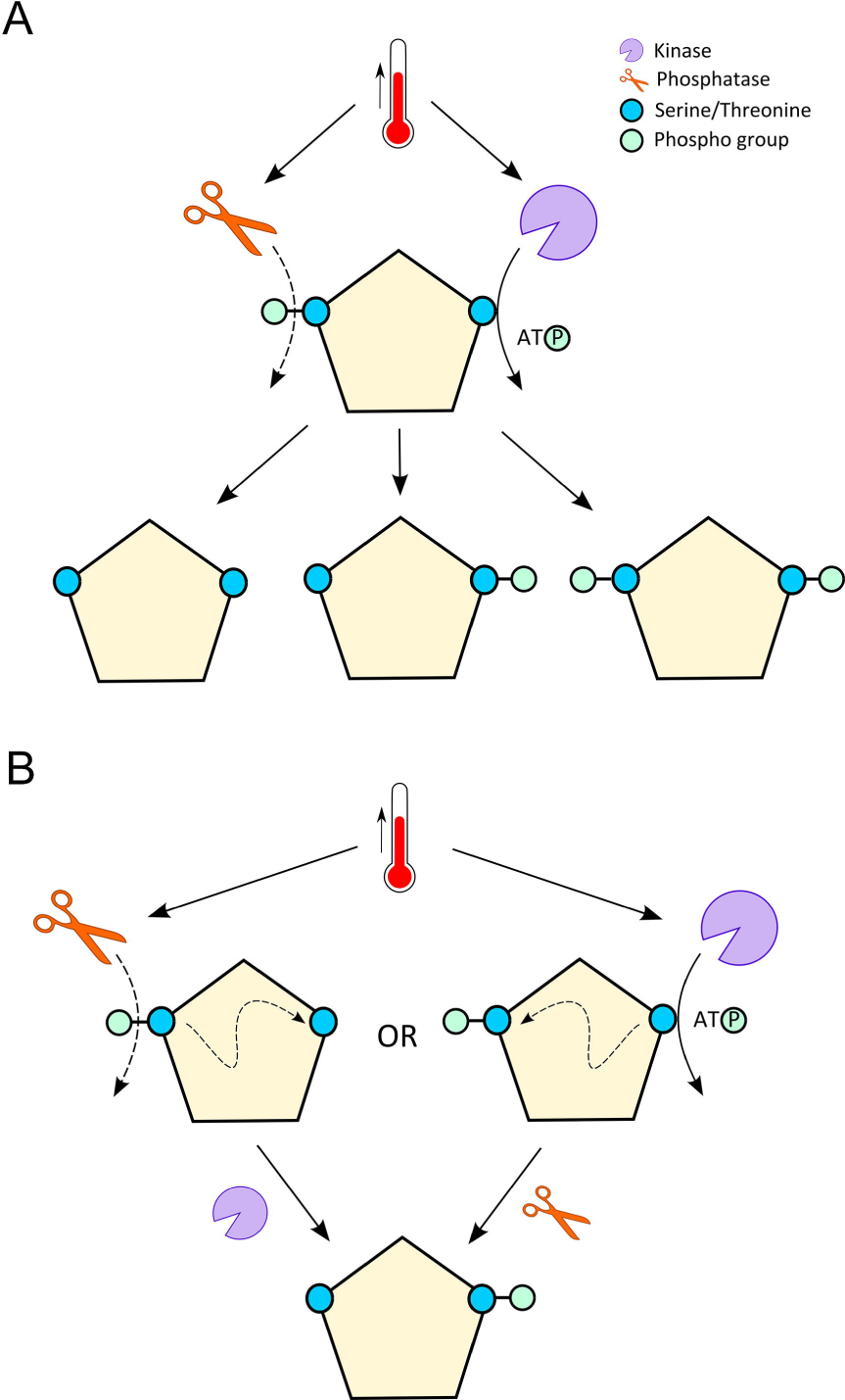
Heat-dependent phosphorylation and dephosphorylation on a single target protein. **(A)** Heat activates both the kinase and the phosphatase to target different Ser or Thr residues simultaneously, generating different phosphoforms of the protein. **(B)** First, heat activates the phosphatase or kinase. The dephosphorylation or phosphorylation of the protein serves as a crosstalk signal for a second kinase or phosphatase to operate, generating one single phosphoform of the protein.

**Table 1.**
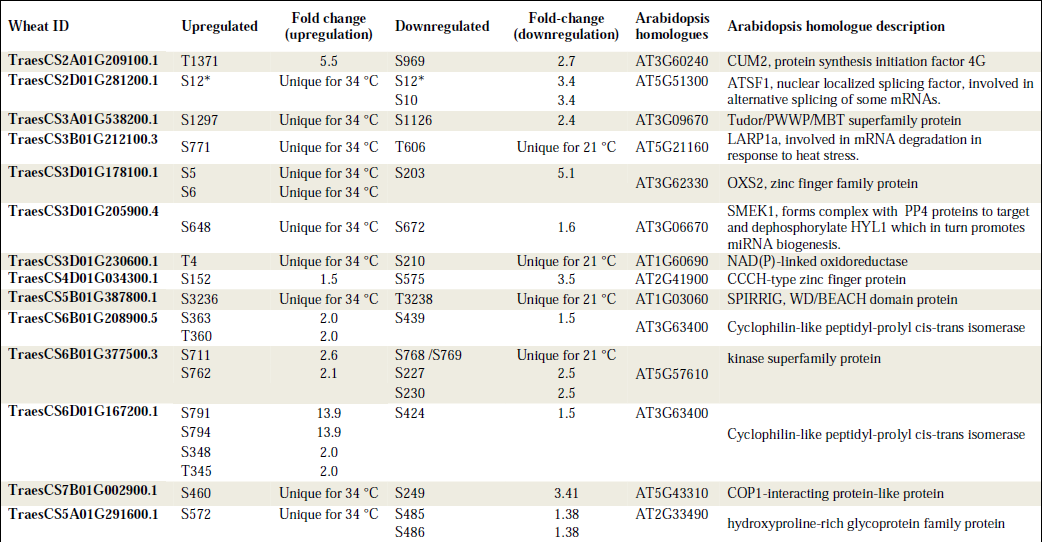
List of phosphoproteins exhibit multiple upregulated and downregulated phosphosites. For TraesCS2D01G281200.1, the peptide containing only phosphorylated S12 is upregulated and the doubly phosphorylated peptide (S12 and S10) is down regulated.

A complex example is the putative protein kinase TraesCS6B01G377500.3 (**Table 1**), which exhibited two phosphosites S711 and S762 that are, respectively, 2.6- and 2.1-fold upregulated in the spikelet samples treated at 34°C. In contrast, a doubly phosphorylated peptide (DFPI***pS***PS***pS***AR, S227 and S230) was detected 2.5-fold higher in the 21°C samples. Further, a single peptide (***pS***SGIETTPAEAEALSK or S***pS***GIETTPAEAEALSK) could only be detected for all 21°C samples, albeit the phosphosite could not be exactly localized (either S768 or S769).

In addition, we also found proteins with multiple phosphosites that showed the same deregulation across different temperature (**Supplementary Table S10**). A large portion of these sites are detected together on the multi-phosphorylated peptides. These phosphosites may work synergistically to control the protein function at elevated temperature or may generate a phosphorylation code for crosstalk between different protein kinases or phosphatases as discussed above. However, in this case, a change in protein level may result in a general change in abundance of phosphopeptide pool. Hence, studying the co-regulation of these phosphosites will require additional investigation on the abundance of the proteins, e.g. by analysing intact proteins or rather the different proteoforms.

Altogether, our data indicated that multiple phosphorylation/dephosphorylation events of a single protein induced by stress are common and add another level of complexity to our understanding of stress signalling mechanisms in plants.

#### Temperature-induced interconversion of neighbouring phosphorylation residues

Interestingly, in the spikelet samples, TraesCS5B01G387800.1 (**Table 1**), which is a homologue of the WD40/BEACH domain protein SPIRRIG in *Arabidopsis thaliana*, exhibited two phosphosites in close proximity with opposite differential regulation upon high temperature. The phosphosite S3236 (***pS***PTTTYGGPGLDVQTLEYR) could only be detected at 34°C, whereas the phosphosite T3238 (SP*p**T***TTYGGPGLDVQTLEYR) could only be detected at 21°C (**Supplementary Figure 4**). The phosphosites are located in the WD40-repeat domain (**Figure 8A**), which is crucial for interaction of SPIRRIG with the decapping protein DCP1 to regulate mRNA decay upon salt stress in *Arabidopsis* (Steffens *et al.*, 2015).

**Figure 8.**
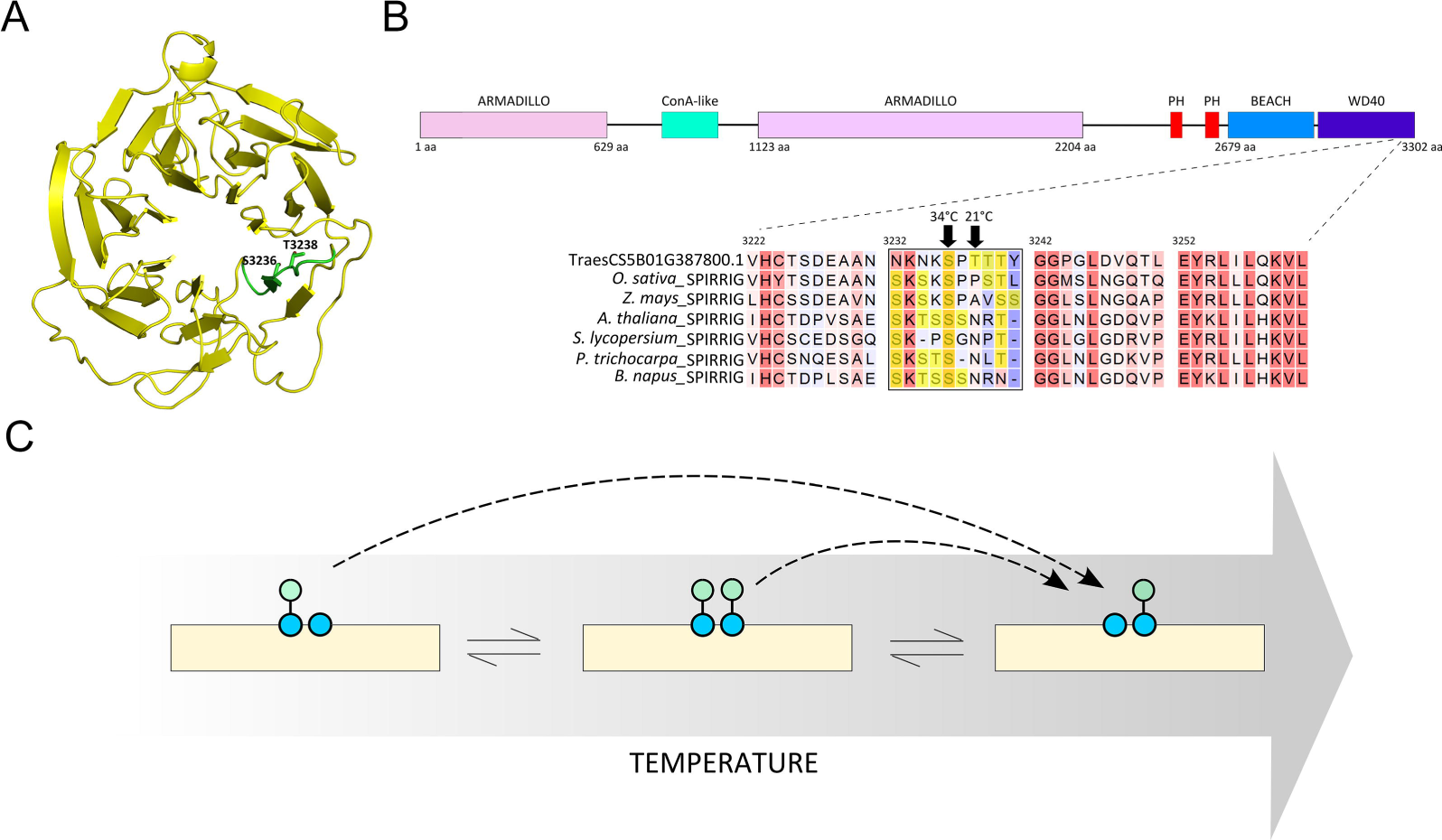
**(A)** Structural model of the WD40 domain of Triticum aestivum SPIRRIG (TraesCS5B01G387800.1). The Ser/Thr rich sequence is highlighted in green showing the two phosphosites detected in the study. **(B)** Alignment of SPIRRIG homologues from different plant species. The Ser/Thr-rich window is marked with the Ser/Thr residues highlighted in yellow. Domain prediction was performed in Interpro (www.ebi.ac.uk/interpro/). **(C)** Model of temperature-induced interconversion of neighboring phosphosites.

Inspecting the protein sequence, we found the two phosphorylation sites localized in a sequence window of 10 amino acids of which four are either Ser or Thr (**Figure 8B**). Neither phosphorylation of the two other Thr residues or a hyper-phosphorylated species of the same peptide could be detected. Hence, a combined effect of phosphorylation of individual sites is likely not relevant. While S3236 was conserved and T3238 not conserved among SPIRRIG homologues, we could also find high frequency of Ser and Thr residues in the same sequence windows in other seed plants (**Figure 8B**). While the high occurrence of phosphorylatable sites might help to preserve the functional phosphorylation pool of a particular sequence during evolution, we suspect that the conformational change of the protein upon stimuli such as heat could lead to the preference for one phosphosite over the other by the same kinase. This might provide a buffering mechanism to maintain the function of the protein by differential phosphorylation of neighbouring amino acid residues depending on the environmental conditions. However, we also do not rule out allosteric or orthosteric regulation between the two phosphosites that might affect the activity of the protein (Nussinov *et al.*, 2012).

For the splicing factor TraesCS2D01G281200.1 (**Table 1**) the phosphorylated peptide containing only S12 (ASAETLARSP***pS***REPSSDPPR) is uniquely detected at 34°C, while the doubly phosphorylated peptide of S10 and S12 (ASAETLAR***pS***P***pS***REPSSDPPR) was 3.4-fold downregulated at the same temperature in the spikelets. We speculate that the phosphoforms of TraesCS2D01G281200.1 may co-exist in a temperature-dependent stoichiometry.

Such interconversion of neighbouring phosphorylation residues (**Figure 8C**) has until now seldom been observed. One example can be found in the cyanobacteria *Synechococcus elongates,* where the circadian clock is controlled by the oscillating phosphorylation equilibrium between a neighbour serine and threonine in the protein kinase KaiC (Rust *et al.*, 2007). This phosphorylation switch between the two residues is modulated by the stoichiometric interaction of KaiC with KaiA and KaiB, in which the pS-KaiC form antagonize KaiA activity, whereas the pT-KaiC form does not. Similarly, a dual phosphorylation switch has been studied in human (Kilisch *et al.*, 2016). To our knowledge, similar phosphorylation modules have not been reported in plants, especially not in the context of stress responses. It is possible that temperature serves as a signalling switch for such a phosphorylation toggle via regulated interaction with at least a protein kinase and/or phosphatase.

## CONCLUSION

In conclusion, we provide the scientific community with the first large scale phosphoproteome in plants under the control of higher ambient temperature across different temperature-sensitive organs. An in-depth analysis showed that the photosynthetic machinery in the leaf is highly responsive to increased temperature, while epigenetic regulation in the spikelets seems to be tightly regulated by higher temperature in a phosphorylation-dependent manner during reproductive development. Furthermore, we observed a core set of common proteins between both leaf and spikelet, suggesting some conserved mechanisms in these organs when responding the higher temperature. Nevertheless, we also observed a large portion of organ-specific regulation. Finally, we exposed a, so far, not reported mechanism of interconversion of neighbouring phosphorylation residues, which likely plays a key role in temperature signalling. Taken together, our data set increases the understanding of temperature signalling in plants.

## Supplementary Data

**Table S1** Primers used in this study

**Table S2** Phosphosites identified in wheat leaves

**Table S3** Phosphosites identified in wheat spikelets

**Table S4** Phosphosites uniquely present at either 24 °C or 34 °C in wheat leaves

**Table S5** Phosphosites significantly deregulated at 34 °C (Students’ t-test p<0.01) in wheat leaves

**Table S6** Phosphosites uniquely present at either 21 °C or 34 °C in wheat spikelets

**Table S7** Phosphosites significantly deregulated at 34 °C (Students’ t-test p<0.01) in wheat spikelets

**Table S8** Phosphosites that is commonly upregulated or downregulated at 34 °C in both leaves and spikelets

**Table S9** Kinases with deregulated phosphosites in this study

**Table S10** List of proteins with multiple upregulated or multiple downregulated phosphosites

**Figure S1** Histograms show normal distribution of Log2 Intensity of quantifiable proteins (proteins present in only one of two temperatures or having at least 2 valid values per temperature) in leaf (A) and spikelet (B)

**Figure S2** Overpresented GO terms for molecular functions among leaf proteins with (A) upregulated or (B) downregulated phosphosites. Fold-changes are indicated.

**Figure S3** Overpresented GO terms for molecular functions among spikelet proteins with (A) upregulated or (B) downregulated phosphosites. Fold-changes are indicated.

**Figure S4** Mass spectrum of phosphopeptides containing S3236 (A) and T3238 (B) in SPIRRIG homologue TraesCS5B01G387800.1.

## Acknowledgements

We thank Michiel Van Bel for assistance in depositing the data in the PTMViewer. We thank Natalia Nikonorova for fruitful discussions on MS data analyses. L.D.V. is the recipient of a VIB International PhD program fellowship. T.Z. is supported by a grant from the Chinese Scholarship Council.

